# Spotiton: New Features and Applications

**DOI:** 10.1101/230151

**Authors:** Venkata P. Dandey, Hui Wei, Zhening Zhang, Yong Zi Tan, Priyamvada Acharya, Edward T. Eng, William J. Rice, Peter A. Kahn, Clinton S. Potter, Bridget Carragher

**Author notes:** For correspondence*: Bridget Carragher (.

## Abstract

We present an update describing new features and applications of Spotiton, a novel instrument for vitrifying samples for cryoEM. We have used Spotiton to prepare several test specimens that can be reconstructed using routine single particle analysis to ~3 Å resolution, indicating that the process has no apparent deleterious effect on the sample integrity. The system is now in routine and continuous use in our lab and has been used to successfully vitrify a wide variety of samples.

## 1. Introduction

We are developing a new instrument (Jain et al., 2012; Razinkov et al., 2016) for vitrifying samples for cryo-electron microscopy (cryoEM). The field of cryoEM has advanced at a breathtaking pace over the past 5 years, but until recently the method used for vitrifying the sample has remained essentially unchanged since it was first developed 30 years ago. The method (Dubochet et al., 1988) consists of applying a small volume (2-3μL) of sample to the surface of an EM grid typically constructed of a metal mesh (Cu, Au, etc.) covered by a support film (typically C or Au), that is often fenestrated. Most of the sample is then wicked away using filter paper to leave a very thin layer (on the order of 10 -100nm) supported by the holey substrate and this is rapidly plunged into a good cryogen (typically liquid ethane or ethane/propane mix). The thin sample is vitrified in this process and can then be transferred into the transmission electron microscope (TEM) where images are acquired of the sample suspended over the holes using low dose conditions. While some instrumentation (e.g. FEI Vitrobot, Gatan CP3, Leica GP) has been developed to make the mechanics of vitrification somewhat easier for novices, some disadvantages of the method remain, i.e. most of the sample is removed by the filter paper and the thickness of the vitrified layer is uneven and sometimes unpredictable.

The instrument that we developed, which we call Spotiton, uses a piezo electric dispensing head to deliver small droplets (10’s of picoliters) to a “self-blotting” nanowire grid. We previously demonstrated that we could vitrify a variety of samples (70S ribosomes, HA, Apoferritin) by using Spotiton to apply small volumes (~2.5-16 nL) of sample to nanowire grids backed by holey carbon substrates. The nanowires effectively act as blotting paper to wick the sample from the surface as soon as the liquid comes into contact with the nanowires, leaving behind a thin wet film that is then plunged into liquid ethane for rapid vitrification.

In this paper we present the current version of this instrument that includes a number of upgrades and improvements. A companion paper provides details about how to manufacture the self-blotting nanowire grids using a variety of formats and materials. We have used Spotiton and the nanowire grids to prepare several test specimens that can be reconstructed to ~3 Å resolution, indicating that the process has no apparent deleterious effect on the sample integrity. The system is now in routine and continuous use in our lab and has been used to successfully vitrify a wide variety of samples.

## 2. Materials and Methods

### 2.0 Spotiton Instrument

For details of the basic design of the Spotiton vitrification instrument, readers are referred to a previous publication (Razinkov et al., 2016). The current layout of the upgraded instrument is illustrated in Figure 1. The device consists of two robotically controlled systems, a dispenser and a plunger, a variety of cameras, a sample station, a humidity control system, and a vitrification bowl. Using the first robot, sample is aspirated into the piezo head, and the formation of suitable droplets is inspected using a camera (see figure 2). The sample head is then rotated into position perpendicular to the grid held in tweezers carried by the second robot. The tweezers are driven towards the cryogen past the piezo head which dispenses a stream of droplets onto the grid forming a strip of liquid (see supplementary videos 1-4) that is rapidly wicked away to a thin film before the sample is plunged into a liquid ethane cup cooled by liquid nitrogen.

**Figure 1:**
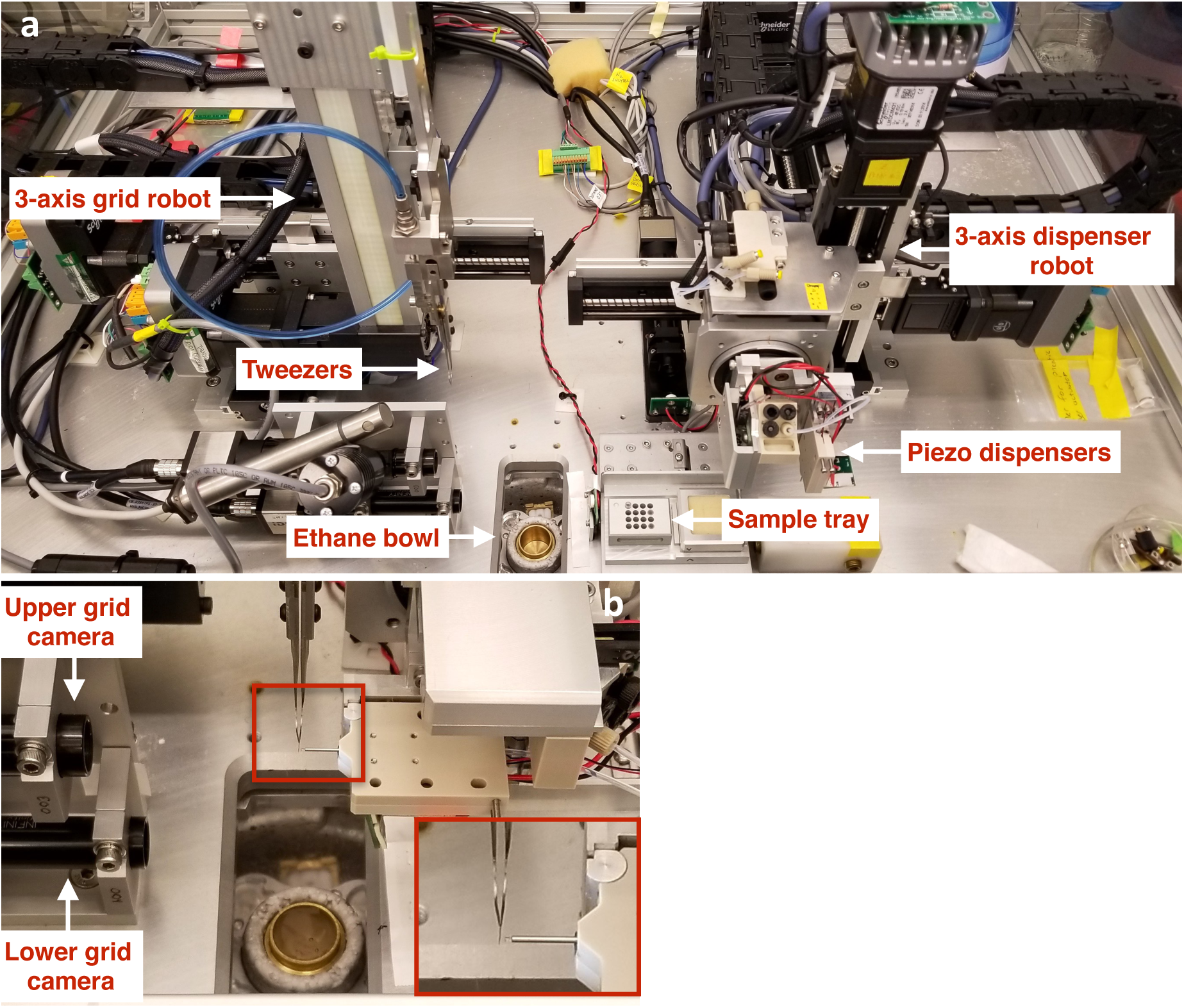
(a) Overall layout of the Spotiton instrument. (b) Grid positioned in front of the piezo tip (magnified view is inset) with upper grid camera viewing dispensing process from behind.

### 2.1 Spotting “on-the-fly”

One of the disadvantages of the previous version of Spotiton was that the time between delivering droplets to the grid and plunging it into the cryogen was too long and varied. Nearly one second elapsed between the time that the dispensed drop hit the grid and the instant when the grid was immersed into the ethane. This dead-time was related to moving the dispense head out of the way (~700ms), opening the shutter between the chamber and the ethane dewar (~100ms), and plunge axis translation (~200ms). As a result, while we had the option to apply drops as small as ~30pL to the grid, the efficiency of the nanowires in absorbing liquid was such that in practice we needed to apply 20-40nL in order to ensure that a wet film remained on the grid at the time of plunging; smaller volume spots would be completely wicked away to leave a dry surface during the travel time it took to immerse the grid in ethane.

In our current version of Spotiton, the droplets are dispensed to the grid as it flies past *en route* to the cryogen. This upgrade required adding optical sensors on the plunge axis to provide position feedback so that dispense heads and cameras can be triggered in precise locations “on-the-fly.” Drops were typically applied using a cosine firing frequency of 14750Hz, corresponding to a distance traveled between spots of 0.338mm. For the results shown in this paper, the plunge speed of the grid was 1m/s with acceleration and deceleration of 10m/s^2^. The time from droplet application to entry into the ethane was ~255ms.

### 2.2 Cameras for optimization

Once the droplets have been applied to the moving grid, we have the option of making a brief stop of the grid in front of a camera to assess the wicking before continuing towards vitrification. The speed of this camera had to be increased to capture the wicking action in more detail. After optimizing back lighting with an LED providing a 4μs strobe pulse we can operate the camera at a speed of ~250frames/s. For the data presented in this paper, the length of time of the stop varied from 1ms to ~120ms, followed by another ~255ms to reach the ethane due to the time of travel. The time interval has to be optimized to ensure that the sample has spread out into a thin film. The rate of dispersal is affected by a variety of factors including the density of nanowires, the dwell time of the grid in the humid environment of the chamber, and the total amount of sample deposited on the grid. A variety of stripes of applied sample viewed by this camera, and subsequently vitrified and imaged in the TEM are shown in supplementary videos V1-V4.

As discussed further below, we would ideally like to minimize the time between sample application and vitrification, and thus we wanted to eliminate or minimize the stop in front of the camera. Initially we expected that the speed of the camera would mean that a strobe capture of a single image would be sufficient to evaluate the sample wicking while the grid flies past the camera. However the strip of liquid applied to the grid cannot be visualized until it begins to wick (see Supplementary videos V1-4) and thus a single frame cannot distinguish between a grid that has not yet started to wick, a grid that is already completely dry, or indeed a failure for the head to fire droplets at all. Evaluating the likelihood of a well vitrified sample is very important at this point of the process so as to avoid the time and effort involved in loading unsuitable grids into the TEM. We thus added a second camera, henceforth referred to as “lower camera”, at a distance of 48mm below the upper camera to grab a snapshot of the grid just before it enters the liquid ethane. The lower grid camera is ~60mm above the ethane bowl which corresponds to ~80ms travel time from this snapshot to vitrification. The upper and lower cameras are aligned to each other and both cameras are supported by two LED strobe lights that can capture a good clear image of a fast moving grid.

### 2.3 Reducing sample volume

One of the other disadvantages of the previous system was that a large dead volume (~30μL) of sample was required for aspiration into the piezo head. We redesigned the sample station to reduce the dead volume so that we now require only 5μL of sample for reliable aspiration, and can use as little as 1μl. Starting with 5μL, usually ~3μL is aspirated into the tip and is used as follows. The first step after aspiration is to test the dispensing of the sample in front of the inspection camera (see figure 2) and we typically use ~30 droplets (~50-75pL/droplet; 1.5nL to 2.2 nL) each time we test. After passing inspection, the tip is positioned in front of the upper grid camera to examine the wicking properties for the specific sample and nanowire grid combination. This step is done by spotting the sample off to one side of the central area of the grid; typically we will repeat this process 3 to 6 times using ~80 droplets (~4nL to 6nL) each time. The goal of these tests is to estimate the optimal conditions so that the wicking will result in a thin wet film at the instant the grid reaches the liquid ethane. The only parameter that is varied is the length of time the nanowire grid is exposed to the humid environment inside the chamber. If we do not wait long enough the grid wicks very efficiently and the sample may be completely wicked away resulting in a dry grid, whereas if we wait too long the wicking efficiency is lowered and the resultant ice will be too thick. Once we are satisfied with the wicking speed we use the same settings to apply the final ~80 droplets now across the center of the grid and the grid is plunged into the liquid ethane. The testing and final sample application consumes ~20nL to ~50nL per grid. The starting volume of 3uL of aspirated sample is thus sufficient to make on the order of ~100 grids. Typically we make between 2 and 8 grids; 2 when we have experience with the sample and understand its properties well enough to be confident of the outcome and several more when working with a new sample.

**Figure 2:**
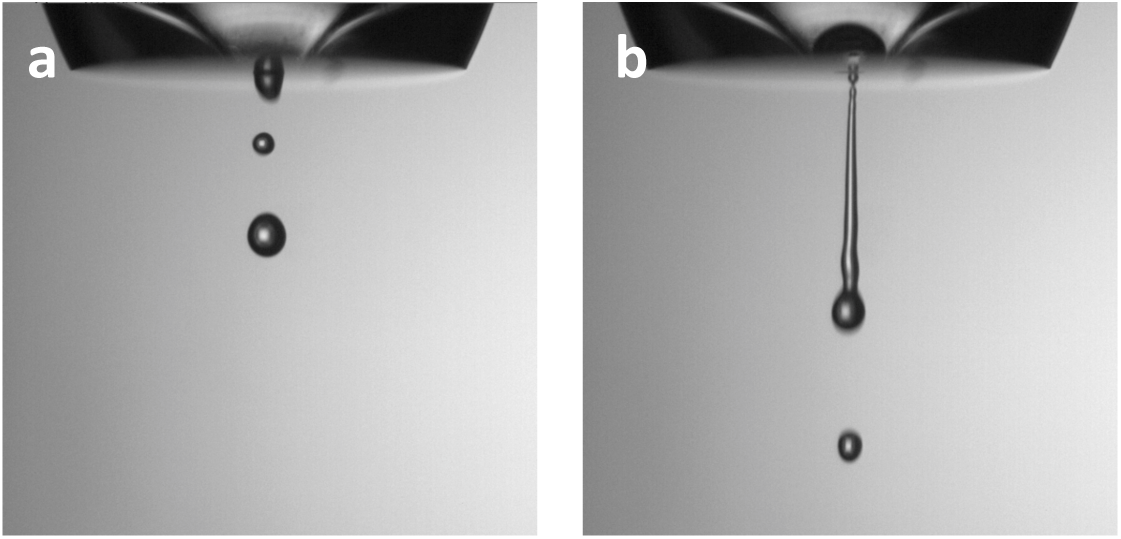
Camera inspection of drop dispensing using cosine mode: (a) satisfactory drop dispensing, (b) unsatisfactory drop dispensing.

### 2.4 Spotiton operation

A typical protocol for operating Spotiton is as follows. On start up, the Spotiton robots are moved to their home position; this procedure includes the two three axis drives supporting sample handling and grid plunging, and the harmonic drive for the piezo tip holder. After homing the robots, the dispensing system is primed, which is achieved by pushing degassed water through the system to clear any air bubbles and fill the empty capillary tube of the tip with water; the tip is cleaned and dried with 100% methanol after every use. The dispensing properties are inspected as described above using water from the reservoir bottle, and once satisfactory, sample is aspirated into the tip and sample dispensing is inspected. The dispense properties are controlled by the mode of operation (cosine or trapezoidal), and frequency and amplitude of the piezo drive pulse. Spotiton is typically operated in cosine mode at a fixed frequency (14750Hz) and the only variable is the amplitude, which relates to the force of the acoustic wave used to break the meniscus of the liquid at the tip, and thus free a droplet. The amplitude required to dispense water droplets is initially adjusted to be somewhere between 375 to 600 (0 to 2047 scale where 2047 corresponds to a maximum of 180 volts peak to peak drive voltage). The amplitude varies from tip to tip but for a specific tip the adjustment is usually within a range of 100. If amplitudes within this range are not able to produce a clean stream of droplets, it is an indication that either the tip function is disrupted by an air bubble or protein debris, or the hydrophobic surface of the front face of the tip has deteriorated such that drops are not cleanly expelled. For the former situation the user needs to re-prime the tip using water or back flush the tip with 100% methanol to remove any debris. Figure 2a shows an example of a tip that is dispensing a satisfactory stream of well separated droplets, whereas figure 2b shows a situation where the droplets are merged into a continuous stream. In the latter case, a well separated droplet stream can usually be achieved by reducing the amplitude applied to the piezo-electric drive. Once a good stream of water droplets is achieved, the sample is aspirated and the amplitude readjusted to achieve good sample dispensing. Again the amplitude will vary according to factors like protein concentration and additives (e.g. detergents and glycerol), but in general the amplitude range is in the region of 400-700, occasionally being set as high as 1200. Other than optimizing the amplitude to deliver well separated droplets, the only other aspect the user has to consider is keeping the surface surrounding the tip orifice dry and hydrophobic.

Once the drop inspection process is completed, the tip is moved in front of the upper grid camera for centering calibration. In this process, the center of the tip is viewed by a video camera and an edge detector algorithm is used to identify the circular curvature of the tip and automatically center it so that the tip can be subsequently aligned to any desired location on the grid for sample dispensing.

The humidity of the chamber is controlled by a humidifier attached to two fans, these fans push the humid air through filters and circulate the air throughout the instrument chamber thus eliminating the formation of airborne large droplets which can settle onto the grids and reduce their wicking capacity. We usually raise the relative humidity of the whole chamber to ~85% once we are ready to start dispensing sample onto the grid. As described above, the sample thickness is optimized by observing wicking behavior using areas near the periphery of the grid. Note that the system has sensors and triggers positioned so as to minimize the volume of the sample dispensed, i.e. the tip is only dispensing sample when the grid is passing in front of it. The first test stripe is applied and the grid is observed using the upper grid camera to estimate the time required to completely wick away all sample (typically 40-200 ms). The user then adjusts the dwell time of the grid inside the ~85% RH chamber so as to partially saturate the nanowires until the wicking time is at least 200ms. This timing is matched to the approximate time of spotting onto the grid and its plunge into ethane (assuming no pause in front of the upper camera). If the wicking time is estimated to be greater than ~200ms the user can define a wait time in front of the upper camera, otherwise the grid proceeds in one continuous motion into the ethane. Typically, 1ms to 300ms delay times were used for the experiments described in this paper. These pause times in front of the upper camera are currently estimated by visual inspection, and we have noted that the estimates may be off by +/-100ms and still produce grids with satisfactory vitreous ice across the entire strip. Following sample dispense, the upper camera acquires a video, if a pause is set, or a snapshot, if the plunge is continuous, followed by a snapshot taken by the lower camera (see figure 3). Various videos of sample wicking acquired using the upper camera are shown in Videos (V1-V4).

**Figure 3:**
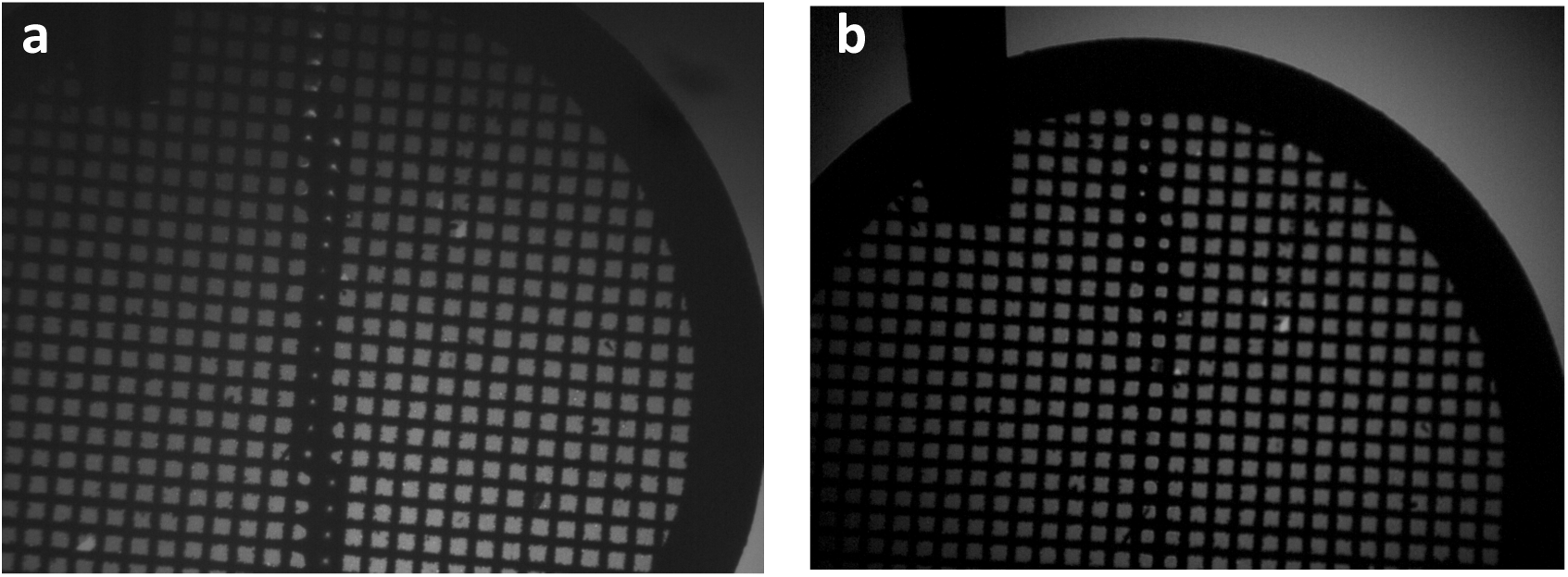
Snapshots of the grid flying past the (a) upper grid camera and (b) lower grid camera after sample deposition.

Following the plunge into ethane the grid is automatically transferred from the ethane into a storage area under liquid nitrogen (see figure 1). The transfer speed of this process was increased as we occasionally observed evidence of devitrified ice that we attributed to the grid warming up during this transfer. The speedup was achieved by encoding the required movements directly into the motors of the three-axis robot which decreased the grid transfer time from ~1000 ms to ~380 ms; note that a manual transfer is typically ~300ms. The grid is then subsequently manually moved to a grid box for storage until imaging.

### 2.5 Imaging and analysis

Data was acquired using Leginon MS (Suloway et al., 2005) on a Titan Krios operated at 300 KeV and equipped with either a K2 direct electron detector or a Bioquantum energy filter and K2 direct detector. MotionCor2(Zheng et al., 2017) was used for drift correcting and dose weighting the movie frames; CTF was estimated using Ctffind4 (Rohou and Grigorieff, 2015). Typically a small set of particles was manually picked, and 2D class averages were calculated using the CL2D algorithm (Sorzano et al., 2010) inside the Appion image processing pipeline (Lander et al., 2009). A subset of these classes were used as templates to pick particles for the entire set of micrographs using FindEM (Roseman, 2004). The stack was then exported to Relion (Scheres, 2012) and taken through a series of 2D and 3D sorting steps before a final 3D refinement was reconstructed using the 3D auto refine option from the Relion-2 package. The directional FSC and histogram plots are generated using 3DFSC program (Lyumkis et al., 2017; Tan et al., 2017).

## 3. Results and Discussion

The upgraded Spotiton instrument is now in regular and routine use in our lab. It has been used to spot dozens of different samples. Samples prepared successfully include small and large soluble proteins, integral membrane proteins, samples with and without detergents, in nanodiscs, and in the presence of glycerol. As discussed above, the only parameter that is varied in making these grids is the amplitude of the acoustic pulse, the wait time inside the humidity chamber and the wait time after spotting. All of these adjustments could be readily automated using image processing algorithms.

We have used Spotiton to reconstruct ~3Å maps (see figures 4-7) of several test specimens (20S proteasome, Apoferrtin), as well as 3-4Å maps of many structures that are the focus of ongoing research projects (see figure 8). These include a 3.4Å map of an integral membrane protein embedded in a nanodisc (see figure 8a) and a 3.6Å structure of an HIV-1 trimer complex (see figure 8b). All of these maps were obtained from a single grid and a single imaging session. The resolution obtained for these structures indicates that the method of preparing grids using Spotiton and nanowire grids is compatible with preservation of the sample sufficient to enable near atomic structural resolution. In table 1 we provide details of the parameters used to prepare the samples and in table 2 some details of the imaging conditions.

**Table 1:** Specimen preparation.

| Sample | MW | Concentraion (mg/ml) | Molarity (μM) | Grid* | Plasma cleaning | Volume spotted | RH/ Temp | Time from spot to ethane (ms) |
| --- | --- | --- | --- | --- | --- | --- | --- | --- |
| 20S Proteasome | 700kD | 4 | 5.7 | CP (~12 nm) | H2, O2 (14sec) | 80x50pl | ~85% | ~355 (100ms delay) |
| Apo ferritin | 480kD | 5 | 10.4 | CP (~12 nm) | H2, O2 (14sec) | 80x50pl | ~80% | ~ 355 (100ms delay) |
| HIV trimer complex + detergent | 660kD | 0.9(complex)<br>2(fab) | 1.4<br>3.0 | CL (~12 nm) | H2, O2 (14sec) | 80x50pl | ~78% | ~255 (1ms delay) |
| IMP in nanodisc | 225kD | 0.7 | 3.1 | CP (~12 nm) | H2, O2 (14sec) | 80x50pl | ~83% | ~ 375 (120ms delay) |

All samples used cosine mode spotting.
CP – Carbon patterned substrate ; CL- Carbon lacey substrate

**Table 2:**
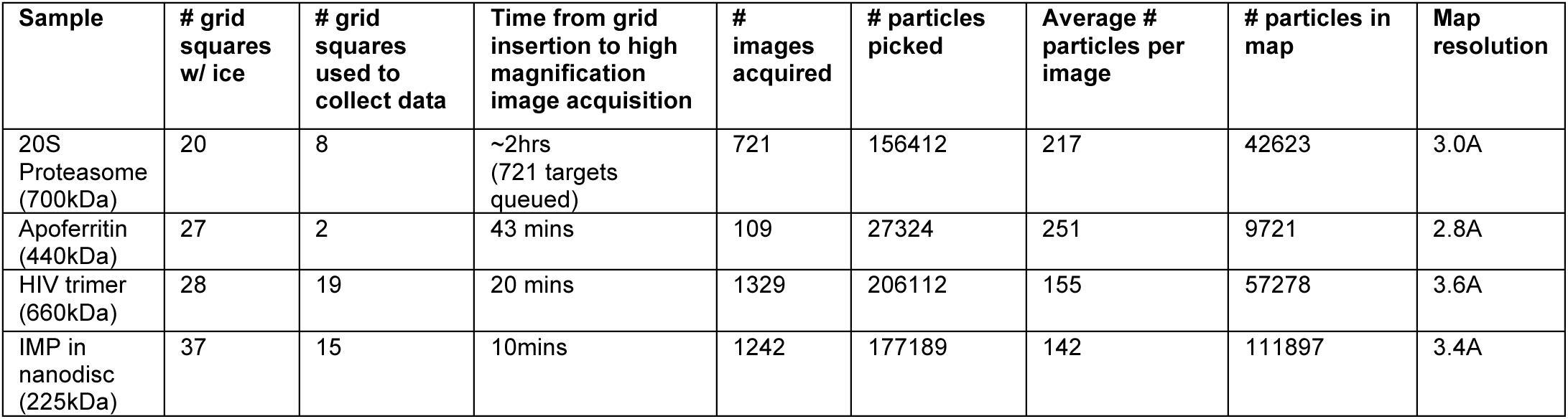
**Imaging and Analysis**

| Sample | # grid squares w/ ice | # grid squares used to collect data | Time from grid insertion to high magnification image acquisition | # images acquired | # particles picked | Average # particles per image | # particles in map | Map resolution |
| --- | --- | --- | --- | --- | --- | --- | --- | --- |
| 20S Proteasome (700kDa) | 20 | 8 | ~2hrs (721 targets queued) | 721 | 156412 | 217 | 42623 | 3.0A |
| Apo ferritin (440kDa) | 27 | 2 | 43 mins | 109 | 27324 | 251 | 9721 | 2.8A |
| HIV trimer (660kDa) | 28 | 19 | 20 mins | 1329 | 206112 | 155 | 57278 | 3.6A |
| IMP in nanodisc (225kDa) | 37 | 15 | 10mins | 1242 | 177189 | 142 | 111897 | 3.4A |

**Figure 4:**
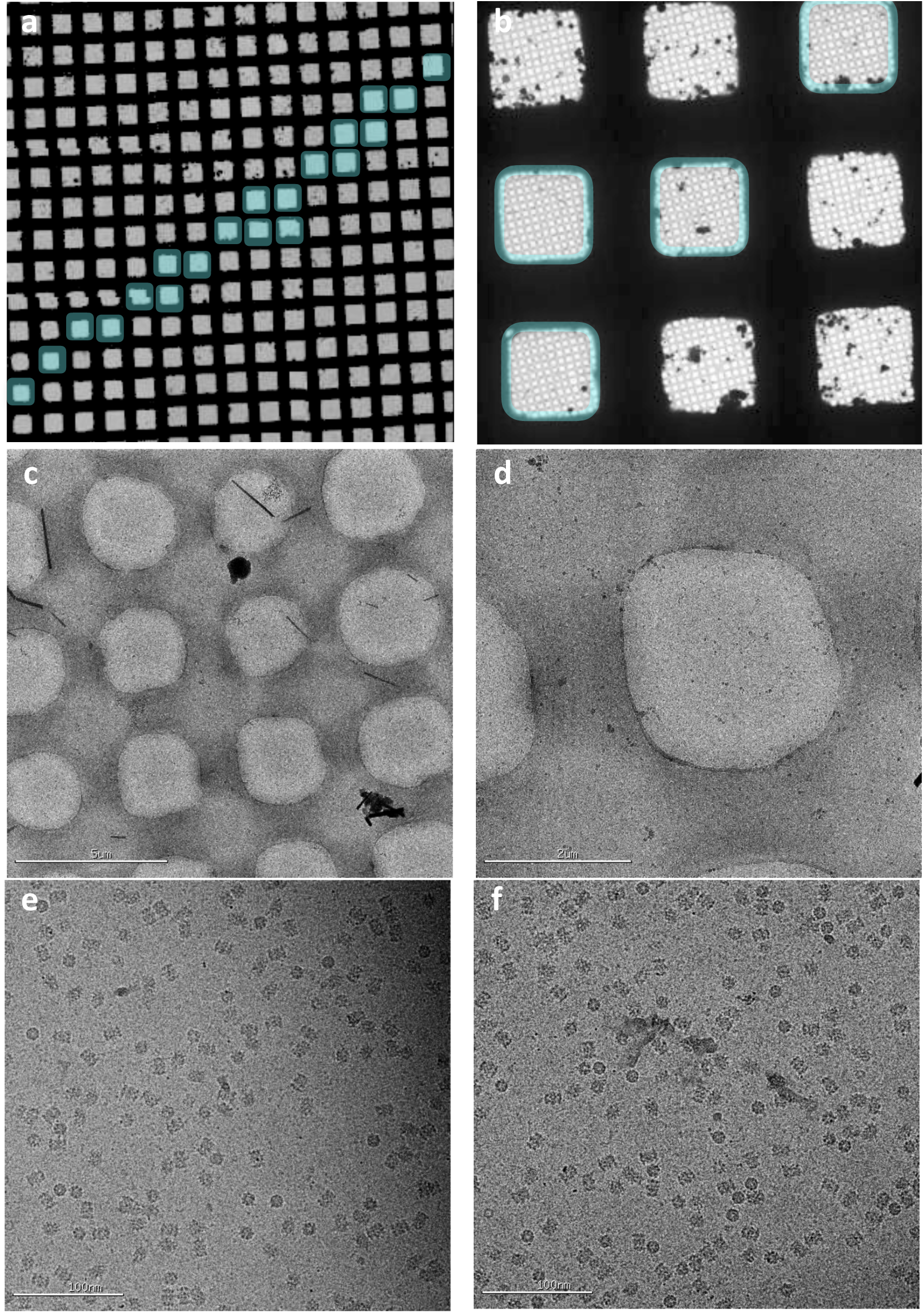
20S proteasome test sample. (a, b) Overall atlas of the grid showing the stripe of vitrified sample shaded blue. (c-d) Sequentially higher magnification images of the vitrified sample. (e-f) High magnification images taken from the hole shown in (d).

A typical data acquisition workflow with Spotiton grids is as follows. Routine alignments of the microscope are performed using the cross-grating grid, the sample grid is then inserted into the microscope, eucentric height is adjusted (5 mins), and an overall atlas of the grid is taken (see figure 4 and 5). The user locates the stripe of vitreous ice across the grid and targets the squares covered with vitreous ice using the multi scale imaging features in Leginon. Because the vitreous ice made using Spotiton is usually uniform and consistent, the time spent on screening the grid to find areas with ice of suitable thickness is much reduced relative to the time to screen grids made using typical protocols. For example, for the 20S proteasome sample (figure 4), after the atlas was completed the user queued up 721 targets in two hours and all imaging was completed within ~11 hours of grid insertion. Similarly, for the Apoferritin sample (figure 5), a queue of 133 exposure targets was set up within ~40 minutes of the atlas imaging and the entire dataset was collected in a period of ~7 hours. The longer imaging time for the Apoferrtin sample was due to super resolution data collection with stage shift which requires ~3 mins per image rather than ~1-2 mins when using counting mode and image shift.

**Figure 5:**
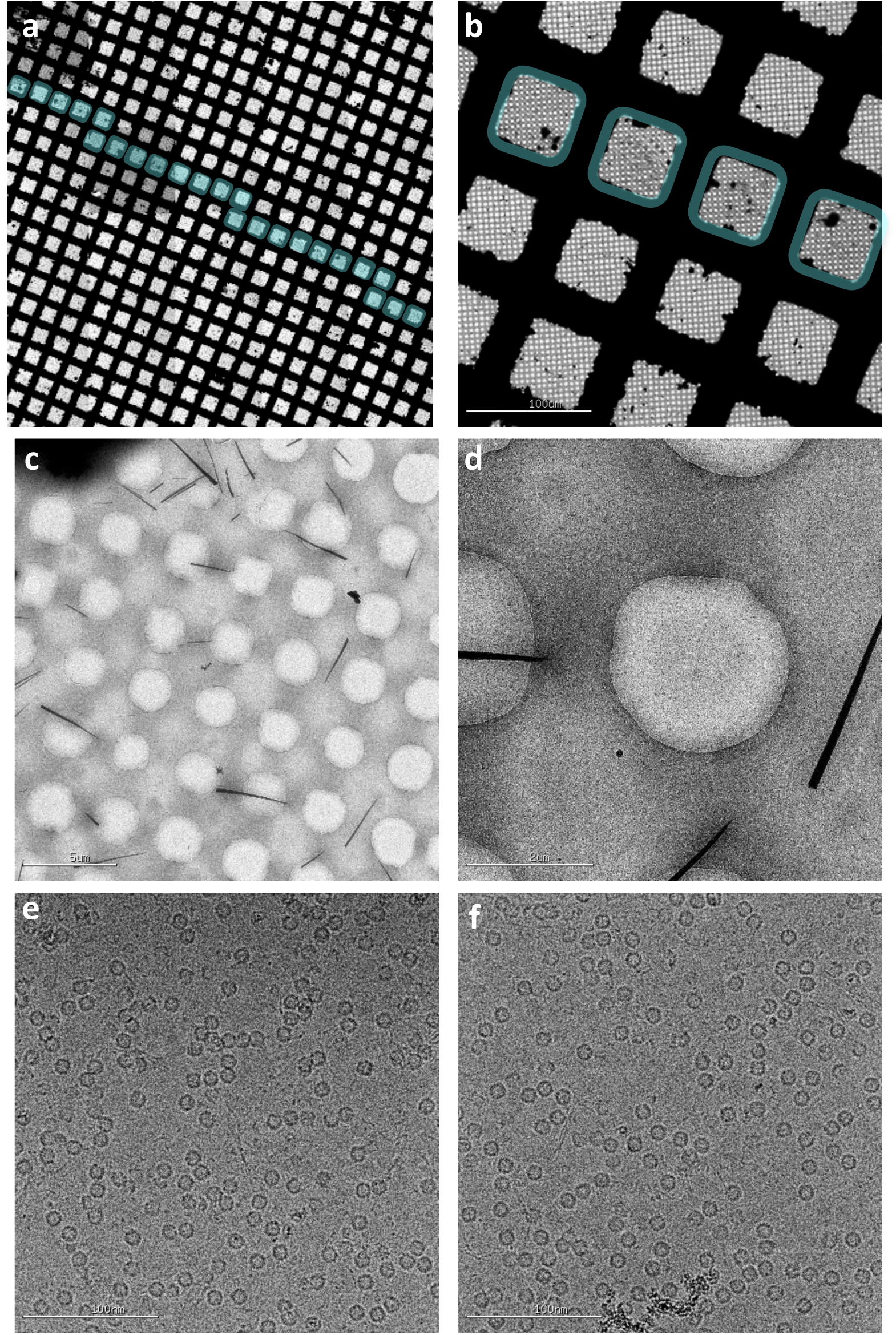
Apoferritin test sample. (a, b) Overall atlas of the grid showing the stripe of vitrified sample shaded blue. (c-d) Sequentially higher magnification images of the vitrified sample. (e-f) High magnification images taken from the hole shown in (d).

**Figure 6:**
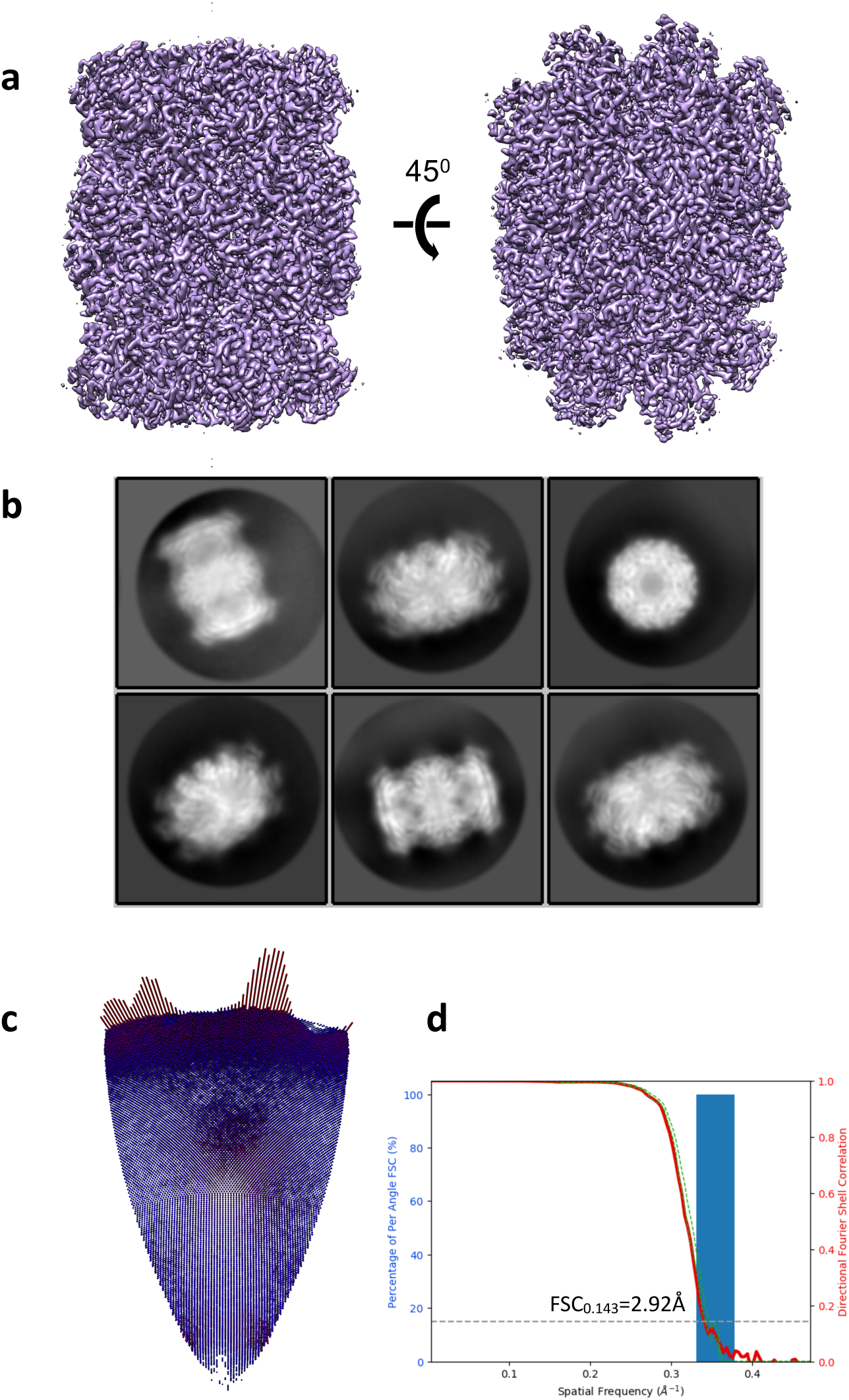
3D reconstruction of 20S proteasome test sample. (a) 3D map rendered with Chimera. (b) Representative 2D class averages. (c) Euler angle distribution. (d) Histogram of directional FSC (blue), global FSC(red) and ± 1 S.D from mean of the directional FSC. FSC curve indicates a resolution of 3.0Å.

**Figure 7:**
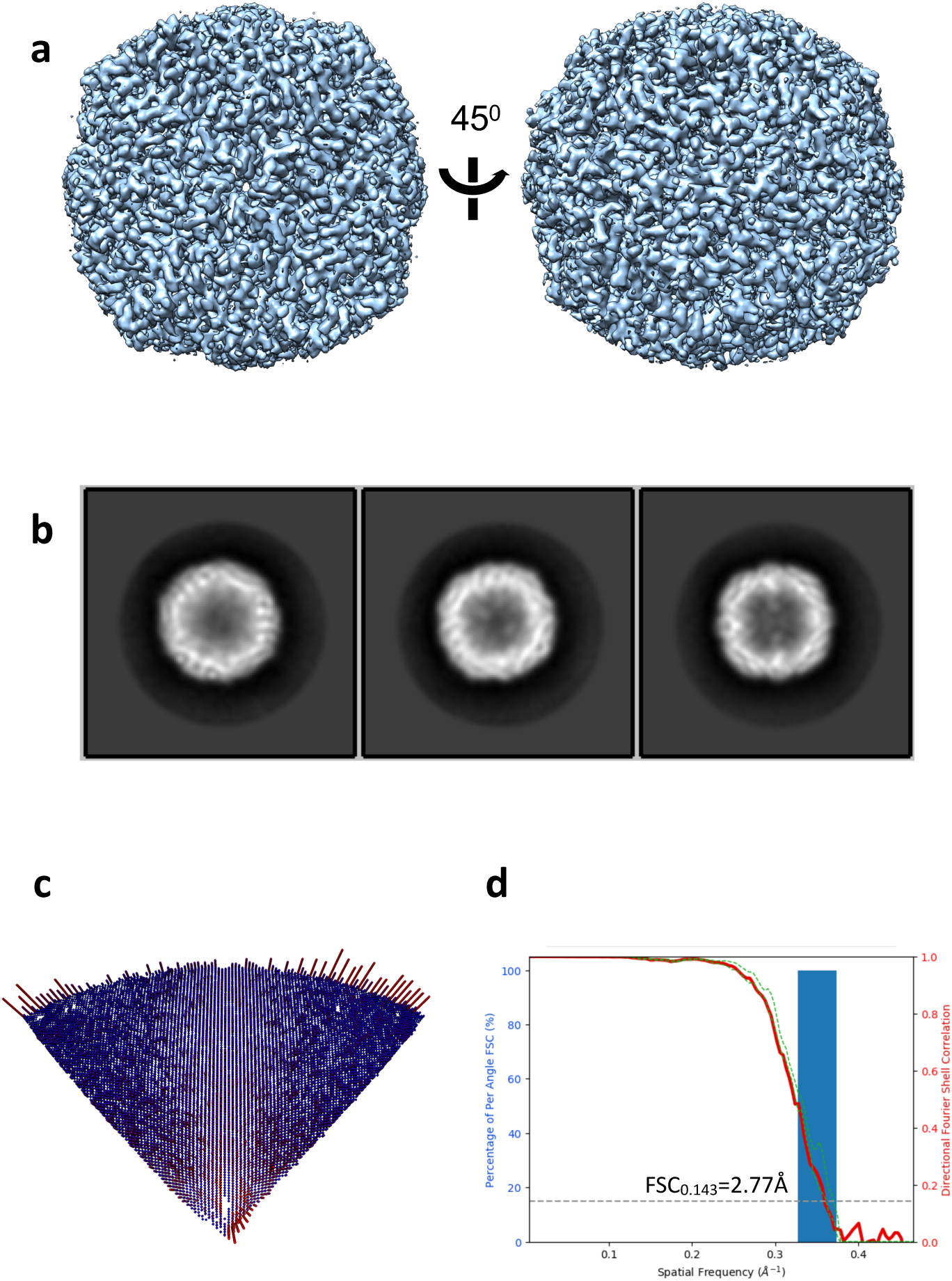
3D reconstruction of Apoferritin test sample. (a) 3D map rendered with Chimera. (b) Representative 2D class averages. (c) Euler angle distribution. (d) Histogram of directional FSC (blue), global FSC(red) and ± 1 S.D from mean of the directional FSC. FSC curve indicates a resolution of 2.8Å.

**Figure 8:**
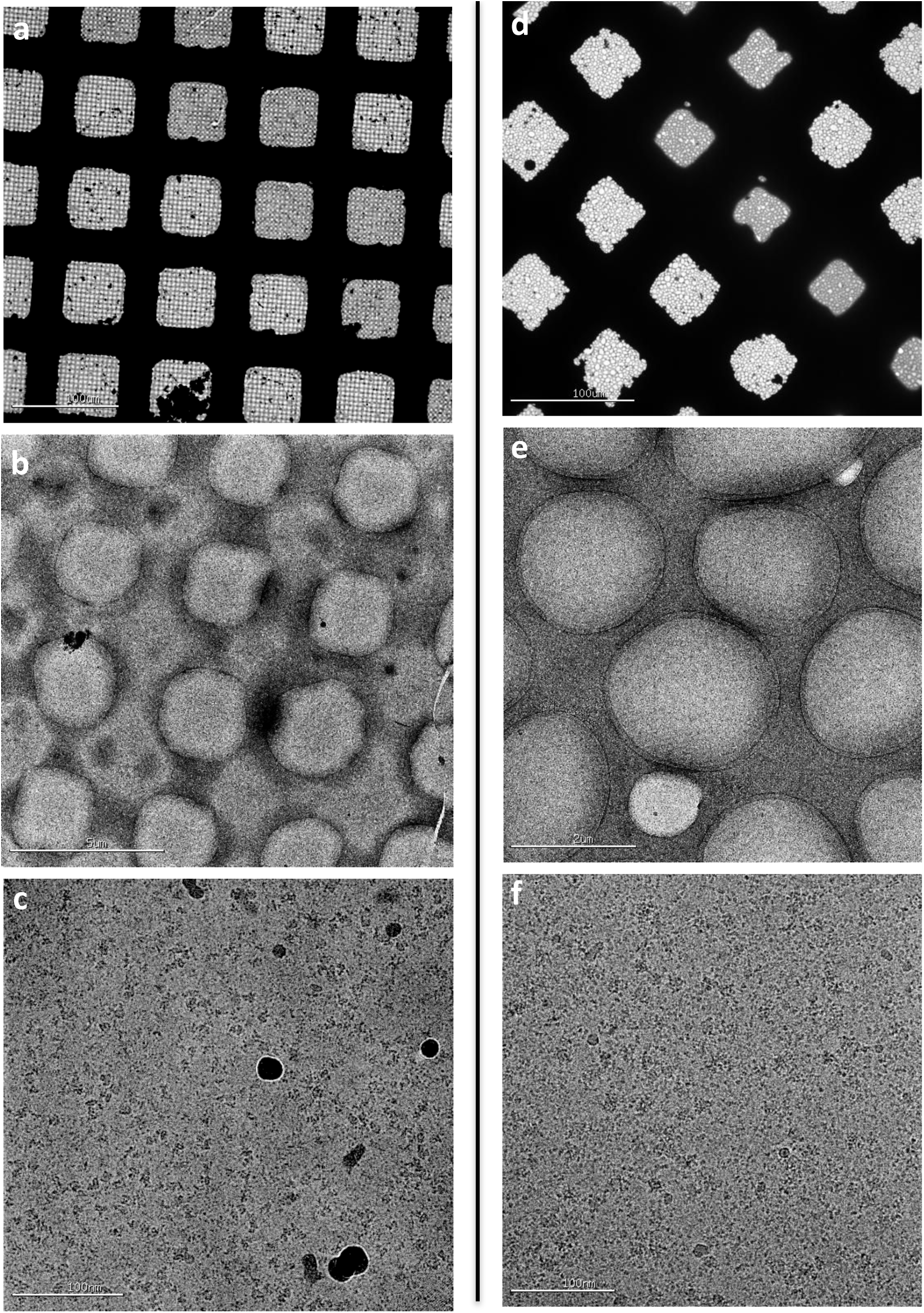
(a-c) Sequentially higher magnification images of a vitrified sample of an integral membrane protein in a nanodisc. (d-f) Sequentially higher magnification images of a vitrified sample of an HIV trimer prepared in the presence of detergent.

There are several areas for further improvements in the technology supporting Spotiton. One of the remaining issues with grid preparation is that particles almost always adhere to the air-water interface and many will be denatured at this interface (Taylor and Glaeser, 2008). This issue is an inevitable consequence of the fact that we must spread our particles out into a very thin layer of liquid prior to vitrification, providing the particles with ample opportunities for colliding, and possibly sticking, to the air water interface. Glaeser (Glaeser and Han, 2017) has postulated that every protein sample will likely denature at the air water interface and the particles we see in vitreous ice are those that are protected by a layer of denatured protein coating this interface. It is certainly true that preferred orientation is a ubiquitous issue for cryoEM and this is likely the result of particles lining up at the air-water interface. Indeed we have observed using tomography, that for ~50 different samples, every one shows particles adhering to one or both of the air water interfaces in the thin layer of vitreous ice (paper under review). Further more,many particles that are perfectly well behaved in negative stain will appear clumped or aggregated in vitreous ice or will disappear completely, presumably because of partial or complete denaturation. These issues are one of the most difficult remaining barriers for cryoEM as a routine and efficient method for protein structure determination. There are several ways to overcome this issue including using a substrate on which to attach the particle and keep them away from the air water interface. Another possibility is to try and “outrun” the issue by limiting the time that the particles are exposed to the air water interface and Spotiton is a good candidate for providing this option. We are working on further shortening the spot to plunge time towards this goal and have some initial indications that this may ameliorate the preferred orientation issue.

Additional areas for improvements include developing image processing algorithms to automate the various adjustments that are made during the grid preparation as discussed above. These steps could include automatically adjusting the amplitude to optimize the spot formation or automatically initiating a cleaning or priming step. We could also envisage an algorithm that could observe the wicking and drying videos and more accurately estimate the the optimal wait time inside the humidity chamber. Our goal in this area is to make the entire process robust and routine and ultimately completely automated. We also anticipate that an automated software controlled system will be capable of more finely estimating the ice thickness so that in the future we should expect to be able to specify the ice thickness range desired for a particular specimen.

## Acknowledgements

We are grateful to Huilin Li and his group (Van Andel Research Institute, Grand Rapids, Michigan) for generously providing the 20S proteasome sample and the staff of the Simons Electron Microscopy Center at the New York Structural Biology Center for help and technical support. The work presented here was conducted at the National Resource for Automated Molecular Microscopy located at the New York Structural Biology Center, supported by grants from the NIH (GM103310, OD019994) and the Simons Foundation (349247).

## 4 Supplementary Information

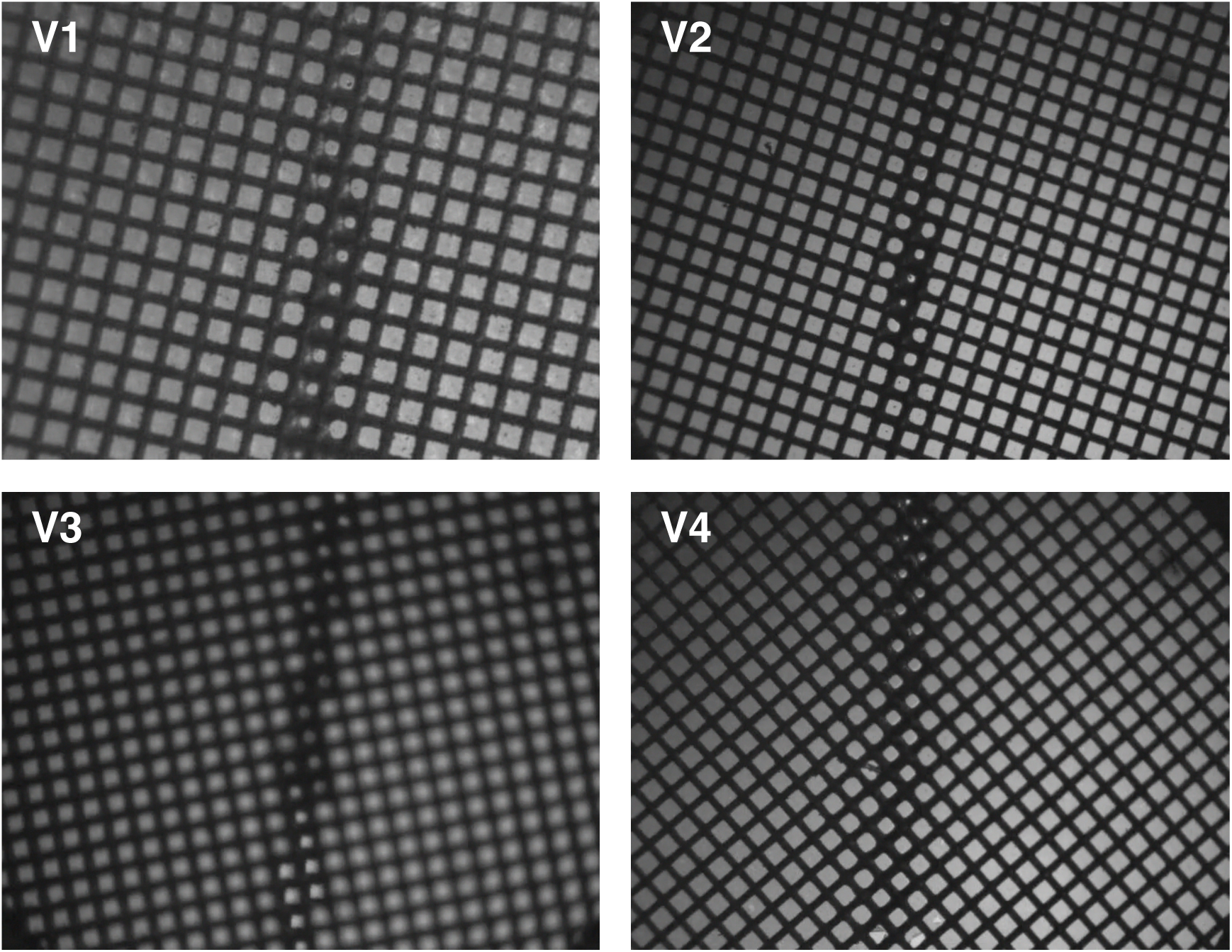
Supplementary Videos of spotting just prior to vitrification:(V1)Apoferrtin;(V2)Proteasome;(V3)HIV trimer;(V4)Integral membrane protein in a nanodisc.

## References

Dubochet, J., Adrian, M., Chang, J.J., Homo, J.C., Lepault, J., McDowall, A.W., Schultz, P., 1988. Cryo-electron microscopy of vitrified specimens. Q. Rev. Biophys. 21, 129–228.

Glaeser, R.M., Han, B.-G., 2017. Opinion: hazards faced by macromolecules when confined to thin aqueous films. Biophys. Rep. 3, 1–7. https://doi.org/10.1007/s41048-016-0026-3

Jain, T., Sheehan, P., Crum, J., Carragher, B., Potter, C.S., 2012. Spotiton: A prototype for an integrated inkjet dispense and vitrification system for cryo-TEM. J. Struct. Biol. 179, 68–75. https://doi.org/10.1016/jJsb.2012.04.020

Lander, G.C., Stagg, S.M., Voss, N.R., Cheng, A., Fellmann, D., Pulokas, J., Yoshioka, C., Irving, C., Mulder, A., Lau, P.-W., Lyumkis, D., Potter, C.S., Carragher, B., 2009. Appion: an integrated, database-driven pipeline to facilitate EM image processing. J. Struct. Biol. 166, 95–102.

Lyumkis, D., Tan, Y.Z., Baldwin, P., 2017. Collecting and processing single-particle cryo-EM data with tilts. Protoc. Exch. https://doi.org/10.1038/protex.2017.055

Razinkov, I., Dandey, V., Wei, H., Zhang, Z., Melnekoff, D., Rice, W.J., Wigge, C., Potter, C.S., Carragher, B., 2016. A new method for vitrifying samples for cryoEM. J. Struct. Biol. 195, 190–198. https://doi.org/10.1016/jjsb.2016.06.001

Rohou, A., Grigorieff, N., 2015. CTFFIND4: Fast and accurate defocus estimation from electron micrographs. J. Struct. Biol. 192, 216–221. https://doi.org/10.1016/j.jsb.2015.08.008

Roseman, A.M., 2004. FindEM--a fast, efficient program for automatic selection of particles from electron micrographs. J. Struct. Biol. 145, 91–99.

Scheres, S.H.W., 2012. A Bayesian view on cryo-EM structure determination. J. Mol. Biol. 415, 406–418. https://doi.org/10.1016/jJmb.2011.11.010

Sorzano, C.O.S., Bilbao-Castro, J.R., Shkolnisky, Y., Alcorlo, M., Melero, R., Caffarena-Fernández, G., Li, M., Xu, G., Marabini, R., Carazo, J.M., 2010. A clustering approach to multireference alignment of single-particle projections in electron microscopy. J. Struct. Biol. 171, 197–206. https://doi.org/10.1016/j.jsb.2010.03.011

Suloway, C., Pulokas, J., Fellmann, D., Cheng, A., Guerra, F., Quispe, J., Stagg, S., Potter, C.S., Carragher, B., 2005. Automated molecular microscopy: the new Leginon system. J. Struct. Biol. 151, 41–60. https://doi.org/10.1016/j.jsb.2005.03.010

Tan, Y.Z., Baldwin, P.R., Davis, J.H., Williamson, J.R., Potter, C.S., Carragher, B., Lyumkis, D., 2017. Addressing preferred specimen orientation in single-particle cryo-EM through tilting. Nat. Methods 14, 793–796. https://doi.org/10.1038/nmeth.4347

Taylor, K.A., Glaeser, R.M., 2008. Retrospective on the early development of cryoelectron microscopy of macromolecules and a prospective on opportunities for the future. J. Struct. Biol. 163, 214–223. https://doi.org/10.1016/j.jsb.2008.06.004

Zheng, S.Q., Palovcak, E., Armache, J.-P., Verba, K.A., Cheng, Y., Agard, D.A., 2017. MotionCor2: anisotropic correction of beam-induced motion for improved cryoelectron microscopy. Nat. Methods 14, 331. https://doi.org/10.1038/nmeth.4193

